# Determinants of transcription factor regulatory range

**DOI:** 10.1101/582270

**Authors:** Chen-Hao Chen, Rongbin Zheng, Jingyu Fan, Myles Brown, Jun S. Liu, Clifford A. Meyer, X. Shirley Liu

**Author notes:** Correspondence should be addressed to C.A.M. or X.S.L.

## Abstract

To characterize the genomic distances over which transcription factors (TFs) influence gene expression, we examined thousands of TF and histone modification ChIP-seq datasets and thousands of gene expression profiles. A model integrating these data revealed two classes of TF: one with short-range regulatory influence, the other with long-range regulatory influence. The two TF classes also had distinct chromatin-binding preferences and auto-regulatory properties. The regulatory range of a single TF bound within different topologically associating domains (TADs) depended on intrinsic TAD properties such as local gene density and G/C content, but also on the TAD chromatin state in specific cell types. Our results provide evidence that most TFs belong to one of these two functional classes, and that the regulatory range of long-range TFs is chromatin-state dependent. Thus, consideration of TF type, distance-to-target, and chromatin context is likely important in identifying TF regulatory targets and interpreting GWAS and eQTL SNPs.

## Introduction

At the time of their discovery, enhancers capable of regulating genes from locations far from the transcription start site (TSS) were a surprising deviation from previous TSS-proximal notions of transcriptional regulation^1,2^. Although the activity of promoters and enhancers has been shown to be dependent on the binding of transcription factors (TFs), the mechanisms through which TFs regulate gene expression are still not well understood. It is now widely accepted that each gene is regulated by the combined influences of several TFs bound nearby^3^, but it is also likely that each TF bound at a single site commonly influences the regulation of more than one gene^4^. The lack of systematic models that accurately describe many-to-many regulatory interactions for TF binding sites limits the accuracy of target gene inference^5^.

ChIP-seq is a broadly used technique for identifying the genome-wide binding sites of specific TFs^6,7,8,9^. Thousands of these binding-site profiles, or cistromes^10^, have been produced in human and mouse cells and tissues^10^. One important use of cistromes is to identify the TFs that regulate a given gene. However, it is difficult to assign most TF binding events to a gene (or vice versa) because relatively few TF binding sites occur very near a gene or within a gene promoter^10^. In most studies target genes are designated using *ad hoc* methods. The simplest of these approaches assign TF binding sites to the nearest gene or to genes based on arbitrary genomic distance thresholds. More accurate approaches^5^ use soft thresholds and consider the effect of multiple binding sites on a gene, but still use arbitrarily defined parameters and model all genes with the same parameters^5^. More recently, genome-wide chromatin conformation Hi-C maps have revealed that the genome is organized as a hierarchically nested structure. Topologically associating domains (TADs)^11,12^ represent one level of this hierarchy and influence enhancer activity by restricting enhancer-promoter interactions between TADs and by facilitating interactions within TADs^11,12^. However, Hi-C maps detect fewer looping interactions between enhancers and promoters than would be expected based on known regulatory interactions^13^ so HI-C alone does not provide a general solution for regulatory interaction inference. How best to use Hi-C observations in quantitative models of cis-regulatory interactions remains an open question. In principle, systematic analysis of Hi-C data, TF cistromes, histone mark ChIP-seq, and gene expression data could reveal insights into gene regulatory mechanisms that lead to better predictions of TF target genes and, in turn, more accurate functional interpretation of non-coding GWAS hits.

In this study, we systematically modeled the genomic distance over which TFs regulate genes, and evaluated how these regulatory ranges depend on the specific TFs and on the genomic and chromatin context of the TAD (as measured by H3K27ac ChIP-seq^14^). Our integrative analyses of large compendia of ChIP-seq^10^, gene expression^15^, and eQTL^16^ data revealed a previously undescribed relationship between transcription factor regulatory range and local genomic and chromatin context. These results suggest the existence of two distinct classes of transcription factors and corresponding mechanisms of gene regulation, which has implications for the analysis of cistromes and disease-associated SNPs.

## Results

### Different regulatory decay distances define two classes of TF

TF cistromes produced by ChIP-seq represent the locations of TF binding sites and potential cis-regulatory elements. To infer the gene regulatory characteristics of different TFs, we used the previously described regulatory potential (RP)^17^ framework to model the relationship between TF cistromes and gene expression. This model assumes that the effect of each single TF binding site on a gene decays exponentially with its genomic distance from the gene’s transcription start site (TSS) (Fig. 1). The contributions of all binding sites near the gene are taken into account by summing the contributions of individual binding sites. We did not presume the existence or definition of “promoter” or “enhancer” functional categories. The model has a single parameter, the “decay distance” Δ, which characterizes the range of influence of a TF binding site on a nearby gene, represented as the half-life of the decay function. When decay distance values are small, only TF binding sites near the TSS contribute to the regulation of a gene; when the decay distance is larger more distant binding sites contribute. We reasoned that the ideal decay distance would produce a good agreement between target genes calculated from TF binding sites and target genes calculated from correlations with TF expression across different biological conditions (Fig.1; Supplementary Fig. 1a). Therefore, for each TF, we tested a range of regulatory decay distances from 100bp to 2000kb and determined the TF regulatory decay distance (Δ*) as the decay distance that gives the best association between TF targets calculated from TF binding and TF targets calculated from gene expression correlation (Fig.1; Supplementary Fig. 1a).

**Figure 1.**
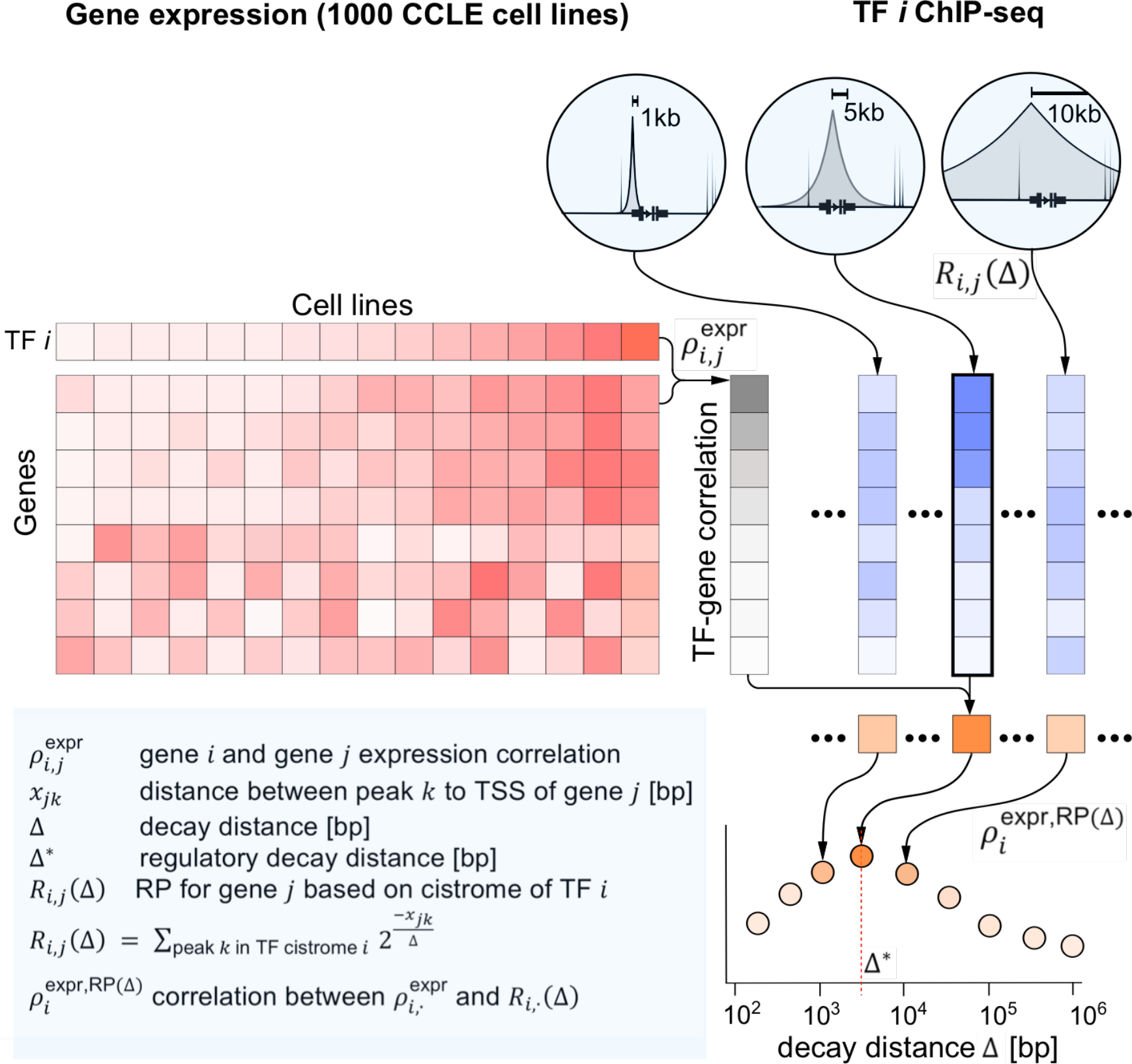
Model for transcription factor influence on gene transcript levels. Schematic representation of the regulatory model. The effect of a single binding site *k* of TF *i* on gene *j* decays exponentially with increasing *x*_*jk*_, defined as the genomic distance between the transcription start site (TSS) of gene *j* and the TF *i* binding site *k*. The decay distance (Δ) is the only parameter in the model and defines the half-life of the exponential decay function 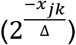. The regulatory potential (RP), *R*_*i,j*_(Δ), defines the total regulatory effect of TF *i* on gene *j* by summing all TF *i* ChIP-seq binding site effects near the gene *j*. To quantify the association between 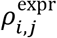 (TF *i* - gene *j* expression correlations using Cancer Cell Line Encyclopedia) and the regulatory potential *R*_*i,j*_(Δ), (see text for rationale) we calculated a second correlation coefficient 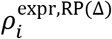 and examined its dependency on the decay distance (Δ). The regulatory decay distance (Δ*) of TF *i* is defined as the decay distance (Δ) that gives rise to the best 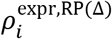 (correlation between *R*_*i,j*_(Δ) and 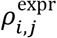).

We first inferred the regulatory decay distances using the TF Cistrome DB collection^10^ and gene expression data across approximately 1000 cancer cell lines from the Cancer Cell Line Encyclopedia (CCLE)^15^. We found that Δ* is less than 1kb for some TFs such as YY1, CREB1, FOXM1, ATF1 and TFDP1 (Fig. 2a, left), but can be greater than 10 kb for other TFs such as PPARG, FOXA1, GRHL2, FOSL2, and TEAD1 (Fig. 2a, right). We thought that ChIP-seq data derived from a limited number of cell lines might bias the inference of regulatory distances, so we repeated the analysis using expression data from GTEx^16^ in distinct tissue types. We observed Δ* values that for a given TF were consistent across tissues (Supplementary Fig. 1b) and that were also consistent with Δ* values calculated with CCLE data. This suggests that our method captures a TF-specific property that persists regardless of the source of the expression data.

**Figure 2.**
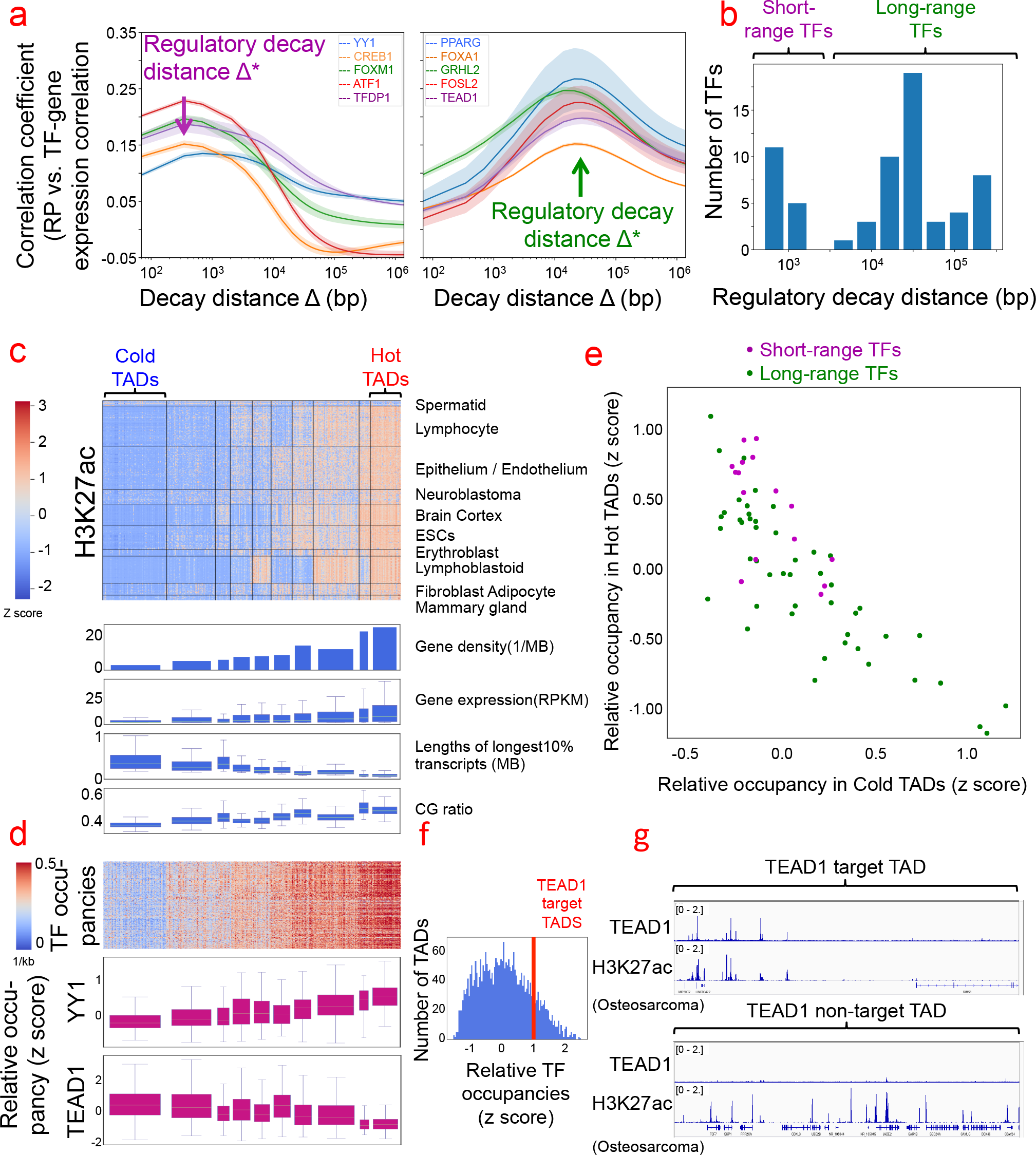
Modeling reveals two distinct TF classes: short-range and long-range. **(a)** Representative TFs with short-range regulatory decay distance (100bp-5kb) include YY1, CREB1, TP53, FOXM1, and ATF1 (left). Representative TFs with long-range regulatory decay distance (5 kb-100 kb) include PPARG, FOXA1, GRHL2, FOSL2, and TEAD1 (right). **(b)** Bimodal distribution of regulatory decay distances of 62 TFs. 16 short-range TFs (left), 48 long-range TFs (right). **(c)** Top: Heatmap of average H3K27ac levels in 3,051 TADs across 1,544 samples from Cistrome DB. 10 TAD clusters (x-axis) and 10 sample clusters (y-axis) were derived by hierarchical clustering. “Cold” and “hot” TADs are indicated. Bottom: Gene densities, average gene expression, lengths of longest 10% transcripts, and G/C fractions correspond to the H3K27ac levels of the TAD clusters. **(d)** Top: Average ChIP-seq peak number per kb in TADs across 3,406 samples with TADs ordered as in (c). Each ChIP-seq sample contains 20,000 peaks. Bottom: the relative occupancies (z-scores) of YY1 and TEAD1 in TAD clusters. Relative occupancy of TF *i* in a given TAD reflects the binding density of the TF in the TAD relative to the binding densities of other TFs in the same TAD. **(e)** Relative TF occupancies in hot and cold TADs. As a trend the short-range TFs have higher relative occupancies than long-range TFs in hot TADs and vice-versa for cold TADs. **(f)** Example of the definition of TF target TADs. TEAD1 target TADs are defined as those TADs with TEAD1 occupancy greater than 1 standard deviation higher than other TFs (Z-scores >=1). **(g)** Illustration of target vs non-target TAD. TEAD1 ChIP-seq and H3K27ac signals in SF269, an osteosarcoma cell line.

Of the 216 TFs with ChIP-seq data in Cistrome DB^10^, we identified 64 TFs with at least three high-quality ChIP-seq samples that yielded high-confidence Δ* estimates (Supplementary Table 1). We observed a bimodal distribution of Δ* (Fig. 2b), suggesting the existence of two distinct classes of TFs: short-range TFs (Δ* between 100bp and 5kb) and long-range TFs (Δ* between 5kb and hundreds of kb). Compared to the cistromes of long-range TFs, those of short-range TFs are significantly enriched within 1 kb of annotated transcription start sites (TSS, Supplementary Fig. 1c) and CpG islands (Supplementary Fig. 1d). This indicates that the distinct regulatory ranges observed in our model recapitulate the biological properties of traditionally defined promoters and enhancers. Because our model is not based on a one-to-one mapping of TF binding sites to the nearest TSS, the result that some TFs act over a short range is not a simple consequence of a TF’s tendency to bind near gene TSSs. Even short-range TFs have a significant proportion of their binding sites outside promoters. Our model suggests that these sites are unlikely to influence gene expression.

### Long-range TFs have unique properties of binding and regulation

Previous studies have suggested that TADs partition the genome into functional domains that help to coordinate gene expression^18,19^. We investigated whether certain TFs or classes of TF may function to regulate the chromatin state and gene expression in designated sets of 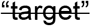 TADs. To this end, we selected 1,545 high-quality human H3K27ac ChIP-seq samples across diverse cell types from the Cistrome DB^10^ and examined their signals across 3,051 previously defined TADs^11^, whose boundaries do not vary across cell type. H3K27ac has been shown to be associated with the promoters of expressed genes and with active enhancers^14^. We clustered TADs across the 1545 samples using the mean H3K27ac ChIP-seq signal in each TAD. This revealed coordinated TAD usage in different cell lineages (Fig. 2c, top). Genes within TAD clusters were enriched in gene ontology categories relevant to the tissue clusters^20^ (Supplementary Table 2). We defined a cluster of “hot” TADs that have high H3K27ac ChIP-seq signal across most samples (316 TADs). We also defined a cluster of “cold” TADs that have low H3K27ac ChIP-seq signal across most samples, but may have high H3K27ac signals in a few restricted tissues (649 TADs, Fig 2c, top). Cold TADs have more A/T-rich DNA^21^, lower gene density, longer gene transcripts, and more tissue-restricted gene expression (Fig. 2c, bottom). A histogram of median H3K27ac signal across samples in the TADs shows a bimodal distribution, with the “hot” and “cold” TADs at the extremes (2086 TADs were not classified as either hot or cold; Supplementary Fig. 1e). Although cold TADs have low levels of H3K27ac, some low level of transcription is evident in these TADs across many cell lines.

To evaluate whether TADs influence TF binding, we examined the binding density of 3,406 TF ChIP-seq samples from the Cistrome DB and observed a universally strong preference of TFs to bind to hot TADs (Fig. 2d, top). To determine whether regulation within a given TAD could be dominated by a given specific TF, we calculated a z-score for each TF-TAD pair to reflect the binding density of the TF in the TAD relative to the binding densities of other TFs in the same TAD. Among other observations, we found that certain TFs like YY1 favor hot TADs, while others, such as TEAD1, favor cold TADs (Fig. 2d, bottom). A systematic analysis shows that the short-range TFs tend to have higher z-scores in the hot TADs, while long-range TFs tend to have higher z-scores in cold TADs (Fig. 2e). Although our model was explicitly ignorant of the existence of “promoters” or “enhancers”, these findings are consistent with well-known principles of gene regulation in which lineage-specifying or cell identity genes are subject to more complex long-range regulation, while ubiquitously expressed genes are regulated at closer range. If a given TAD is enriched for the binding of a TF relative to the binding of other TFs (z-score > 1), it is defined as a “target” TAD of that TF (Fig. 2f). In TEAD1 target TAD (Fig. 2g top) there are nearly perfect co-localization of TEAD1 and H3K27ac ChIP-seq peaks, suggesting that TEAD1 is a TF that dominates the activity of its target TAD. Indeed, the GO enrichment of genes within the target TADs correspond to the known functions of their targeting TFs (Supplementary Table 3).

Cold TADs are likely to be in a default heterochromatin state. We reasoned that TF binding within these TADs might require pioneer-like properties, characterized by the capability to facilitate the binding of other TFs at their binding sites^22^. In the CCLE cell lines, we evaluated whether a TF’s DNA motif becomes more enriched in the ChIP-seq peaks of other factors as the TF’s expression increases. For instance, in cell lines with increased TEAD1 expression, the TEAD1 motif becomes more enriched in the ChIP-seq peaks of other factors (Fig. 3a). TFs with such ‘pioneer-factor-like’ properties (Supplementary Table 4) are generally long-range TFs (Fig. 3b, Methods) and include known pioneer factors (FOXA1^22^, GRHL2^23^) and those not previously known to be pioneers (FOSL2) (Supplementary Fig. 2a).

**Figure 3.**
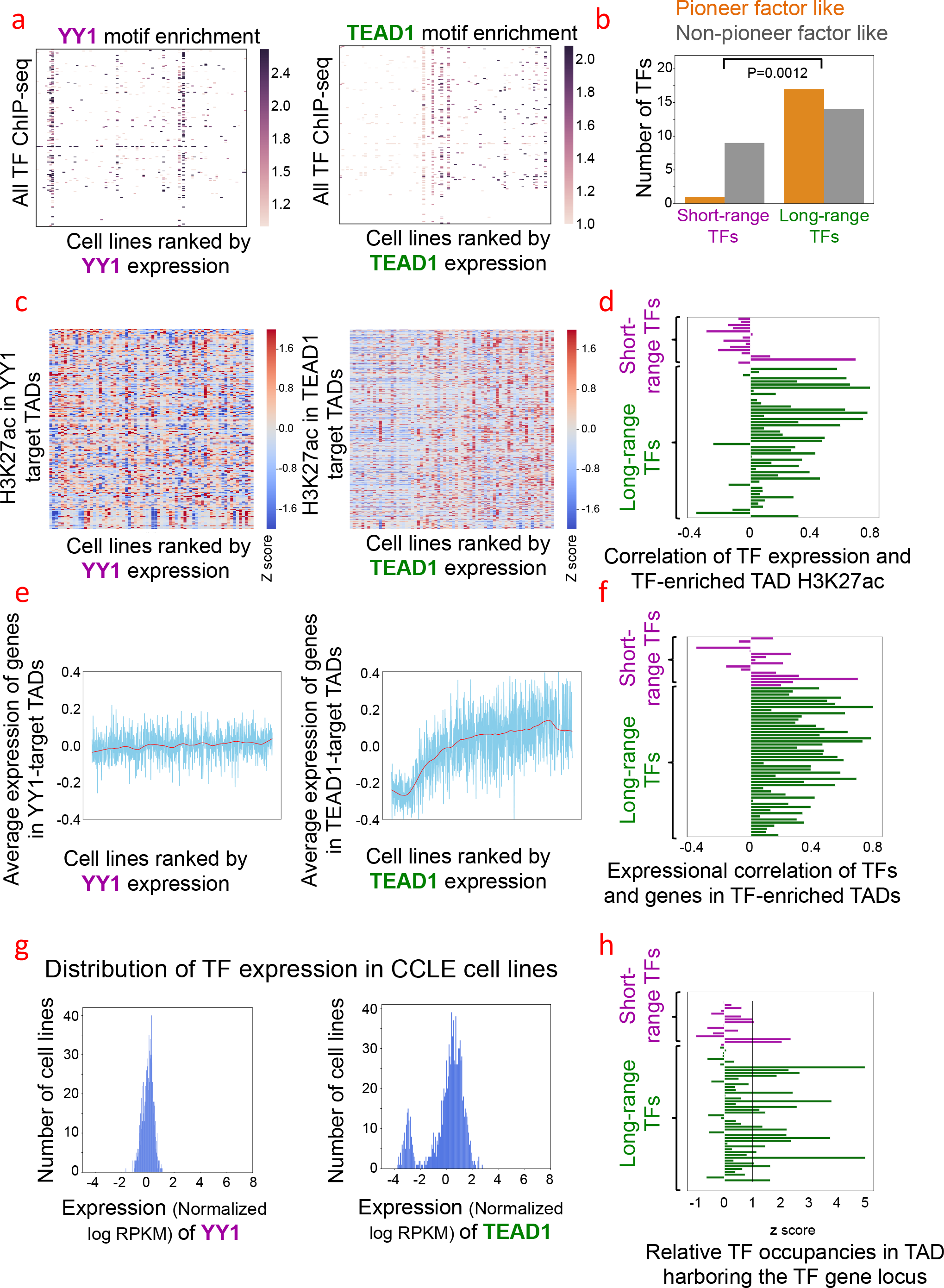
Long-range TFs and short-range TFs have distinct regulatory properties. **(a)** In each panel, the X-axis consists of TF ChIP-seq cell lines ranked lowest to highest by YY1 (left) or TEAD1 (right) expression using CCLE data. The Y-axis in each panel plots TF ChIP-seq samples in Cistrome DB. Each entry represents the enrichment of YY1 or TEAD1 motifs in the ChIP-seq peaks of a TF-cell line pair. No increase of YY1 (a short-range TF) motif enrichment is evident in the ChIP-seq binding sites of factors besides YY1 in cells with high expression of YY1 (left panel), in contrast to TEAD1 (right panel). **(b)** Long-range TFs and short-range TFs with pioneer-like properties. Pioneer-like is defined as a significant increase (logistic regression p-value < 0.01; Methods) of TF motif enrichment in the ChIP-seq binding sites of other TFs in cells with high expression of the candidate pioneer-like TF. **(c)** Left: No increase of H3K27ac levels is evident in YY1 target TADs in cell lines with high YY1 expression. Right: An increase of H3K27ac levels in TEAD1 target TADs is evident in cell lines with high TEAD1 expression (p-value 0.003 using Pearson correlation). X-axis: H3K27ac ChIP-seq cell lines ranked by YY1 (left) or TEAD1 (right) expression using CCLE data. Y-axis: H3K27ac ChIP-seq samples in Cistrome DB. **(d)** Long-range TFs (green) have a stronger correlation between TF expression and H3K27ac levels in target TADs than short-range TFs (magenta). **(e)** Left: The average expression of genes located within YY1 (a short-range TF) target TADs does not increase with increasing YY1 expression. Right: The average expression of genes located within TEAD1 (a long-range TF) target TADs is high in cell lines with high TEAD1 expression. **(f)** Long range-TFs (green) have a stronger correlation between TF expression and the expression of genes located in its target TADs. Short-range TFs (magenta). **(g)** The distribution of TF expression in CCLE cell lines. The expression value (log transformed RPKM) of each gene is normalized by subtracting the mean value across all samples. Left: YY1. Right: TEAD1. **(h)** Long range TFs (green) have multiple binding sites within the TAD that contains the TF gene itself. Short-range TFs (magenta).

To examine whether long-range TFs may influence the chromatin state of target TADs, we examined whether the expression of long-range TFs was robustly associated with H3K27ac levels in the target TADs. For each TF, we selected CCLE cell lines with available H3K27ac ChIP-seq data and sorted these cell lines by the TF expression level. We observed generally positive associations between the TF expression and the mean H3K27ac levels of the target TADs for long-range TFs such as TEAD1 (Fig. 3c, 3d), but weaker or negative associations for short-range TFs, such as YY1 (Fig. 3c, 3d). In addition, there are significant positive associations between the expression of long-range TFs, such as TEAD1, and genes located in their target TADs (Fig 3e, 3f, Supplementary Fig. 2b) and weaker associations for short-range TFs (Fig 3e, 3f, Supplementary Fig. 2b). This suggests that long-range factors have more influence over chromatin state and gene expression in their target TADs than short-range factors.

To evaluate whether the long-range TFs are more likely to be expressed in a lineage-specific manner, we examined the CCLE expression patterns of all TFs with confident decay distance estimates. Long-range TFs, such as TEAD1, tend to have a bimodal or long-tail gene expression distribution across cell lines (Fig. 3g, Supplementary Fig. 2c) in contrast to the unimodal gene expression distribution for most short-range TFs, such as YY1 (Fig. 3g, Supplementary Fig. 2c). Positive feedback loops composed of self-activating TFs have been proposed as a mechanism for establishing stable gene expression programs during lineage specification^24,25^. Indeed, the TSS for 20/48 long-range TFs is located within one of its own TF target TADs, suggesting that long-range TFs might have auto-regulatory properties (Fig. 3h, Supplementary Fig. 2d). This indicates that multiple binding sites of the same TF within the TAD that contains the TF gene itself may serve as a robust auto-regulatory mechanism for maintaining lineage restricted TF expression.

### Regulatory decay distances differ between hot and cold TADs for the same TF

Known cases of gene regulation over distances greater than 100kb occur in TADs that in most cell types have low levels of H3K27ac occupancy^26,27,28^. We therefore hypothesized that enhancers influence genes over longer genomic distances in cold TADs than in hot ones. To test this we first used an independent data type, CAGE-seq^29^, which captures the transcription start sites of both mRNAs and eRNAs. We then compared these density maps between hot and cold TADs and found higher enhancer-promoter correlations over longer genomic distances in cold TADs (Supplementary Fig. 3a). To confirm this finding, we further examined the GTEx eQTL data and found the distances between eQTLs and the corresponding gene TSSs to be significantly longer in cold TADs in most tissue types (Fig. 4a). This phenomenon is not due to linkage disequilibrium (LD) since the LD block sizes are similar in cold and hot TADs (Supplementary Fig. 3b).

**Figure 4.**
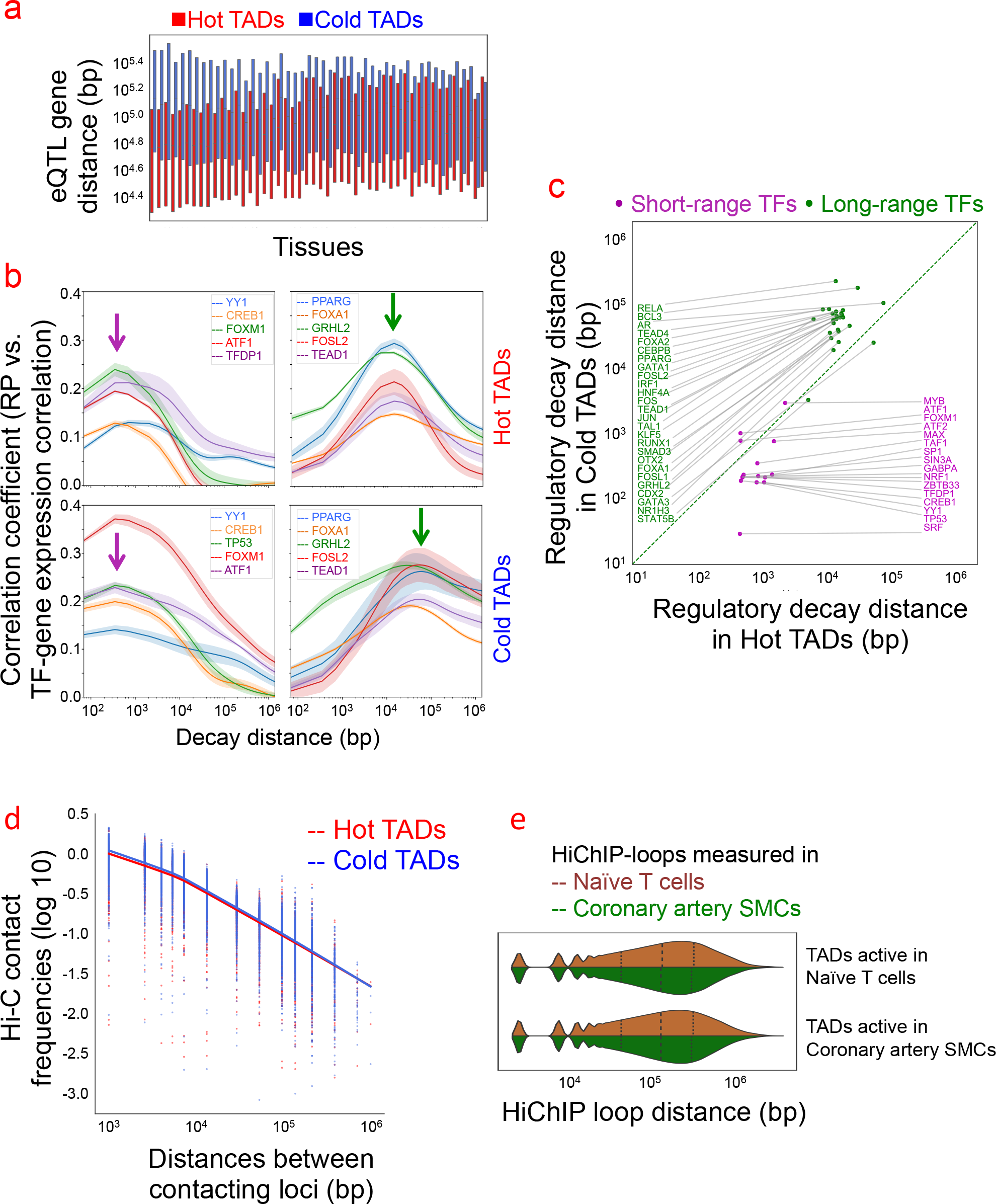
Long-range TFs have longer regulatory decay distances in cold TADs. **(a)** Distribution of GTEx eQTL-TSS distances in hot (red) and cold TADs (blue) across different tissues. In 47 (out of 48) tissues, eQTL-TSS distances are longer in cold TADs than those in hot TADs. TADs were sorted using average eQTL-TSS distances. **(b)** Regulatory decay distance of long-range TFs depends on chromatin context. X-axis: decay distance (Δ); Y-axis: correlation coefficient between RP (*R*_*i,j*_(Δ)) and TF-gene expression correlation 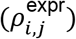. Top row: Hot TADs. Bottom row: Cold TADs. Left column: Short-range TFs, including YY1, CREB1, TP53, FOXM1, and ATF1. Right column: Long-range TFs, including PPARG, FOXA1, GRHL2, FOSL2, and TEAD1. **(c)** Long-range TFs (green dots) have longer regulatory decay distances in cold TADs. **(d)** In hESC Hi-C data, the average contact frequency between two loci decreases as the distances between the loci increases, following a power-law relationship. The decay rates are not significantly different between cold TADs (blue) and hot TADs (magenta). **(e)** Violin plot of H3K27ac Hi-ChIP loop sizes in naïve T cells and coronary artery smooth muscle cells (SMCs). Loops located in TADs more active in naïve T cells (brown) or TADs more active in coronary artery SMCs (green) have similar sizes.

Based on the above observations, we re-modeled TF regulatory decay distances in cold and hot TADs separately, rather than a single global Δ* as we have done in the previous section. Interestingly, we found that for the long-range TFs Δ* becomes even longer in cold TADs (Fig. 4b,c) (relabel 4b). This is consistent with the proposition that the few documented observations of long-range enhancer activity in cold TADs is the norm rather than the exception^26,27,28^. Although TF binding sites and TSSs tend to be sparser in cold than in hot TADs, the difference in decay distances between the cold and hot TADs is not a trivial consequence of the model or the binding site distribution. The regulatory decay distances inferred using the most significant 10,000 and 20,000 peaks are almost the same, suggesting that the model is insensitive to the peak to gene distance distribution (Supplementary Fig. 3c).

We next examined whether the difference in the regulatory behavior in hot and cold TADs is due to differences in chromatin interaction frequencies within these TAD types. We carried out an analysis of Hi-C chromatin interaction data^30^, comparing average contact frequencies within hot TADs to those within cold TADs as a function of genomic distance. Contact probabilities decrease with genomic distance following a power-law relationship^31^, and the decay rates are not significantly different between hot and cold TADs. This suggests that the overall properties of interaction frequencies of hot and cold TADs do not explain the longer regulatory decay distances for long-range TFs in cold TADs (Fig. 4d). We also examined H3K27ac Hi-ChIP data in different cell lines and found that loop distances (Fig. 4e) and contact frequencies (Supplementary Fig. 3d) do not correlate with the H3K27ac level of the TAD. Therefore Hi-C-derived or Hi-ChIP-derived interaction frequencies do not explain the range of regulatory effects, suggesting that either the technology does not capture all important regulatory interactions on the relevant time and length scales, or that mechanisms other than direct physical interactions underlie the long-range regulatory interactions.

### Regulatory decay distances change with chromatin states in the same TADs

Our analyses revealed that the bindings of long-range TFs in hot and cold TADs have different impacts on gene regulation (Fig. 4c). However, several genetic properties of TADs contribute to the establishment of their chromatin states and influence their designation as hot or cold (Fig. 2c). To tease out the true chromatin effect from the genetic properties, we next assessed whether different chromatin states in the same TADs (for example measured in two different cell lines), have different impacts on regulatory distances (Fig. 5a). We identified the nuclear factor NF-kappa-B p65 subunit, RELA, for which gene expression data in GTEx and multiple ChIP-seq datasets are available in the lymphoblastoid cell line GM12878 and lung carcinoma cell line A549. We defined TADs with statistically significant differences in H3K27ac ChIP-seq signals between the two cell lines as GM12878-predominant (more H3K27ac in GM12878 than in A549) or A549-predominant TADs. We then used GTEx lymphoblastoid and lung tissue expression data to find the regulatory decay distance Δ* of RELA by focusing on the GM12878-predominant and A549-predominant TADs, respectively. We observed that RELA Δ* estimated in lymphoblastoid is shorter in GM12878-predominant TADs and the RELA Δ* estimated in lung is shorter in A549-predominant TADs (Fig. 5b). This indicates that the regulatory decay distance for the same TF becomes shorter in the same TADs when the TADs become more active.

**Figure 5.**
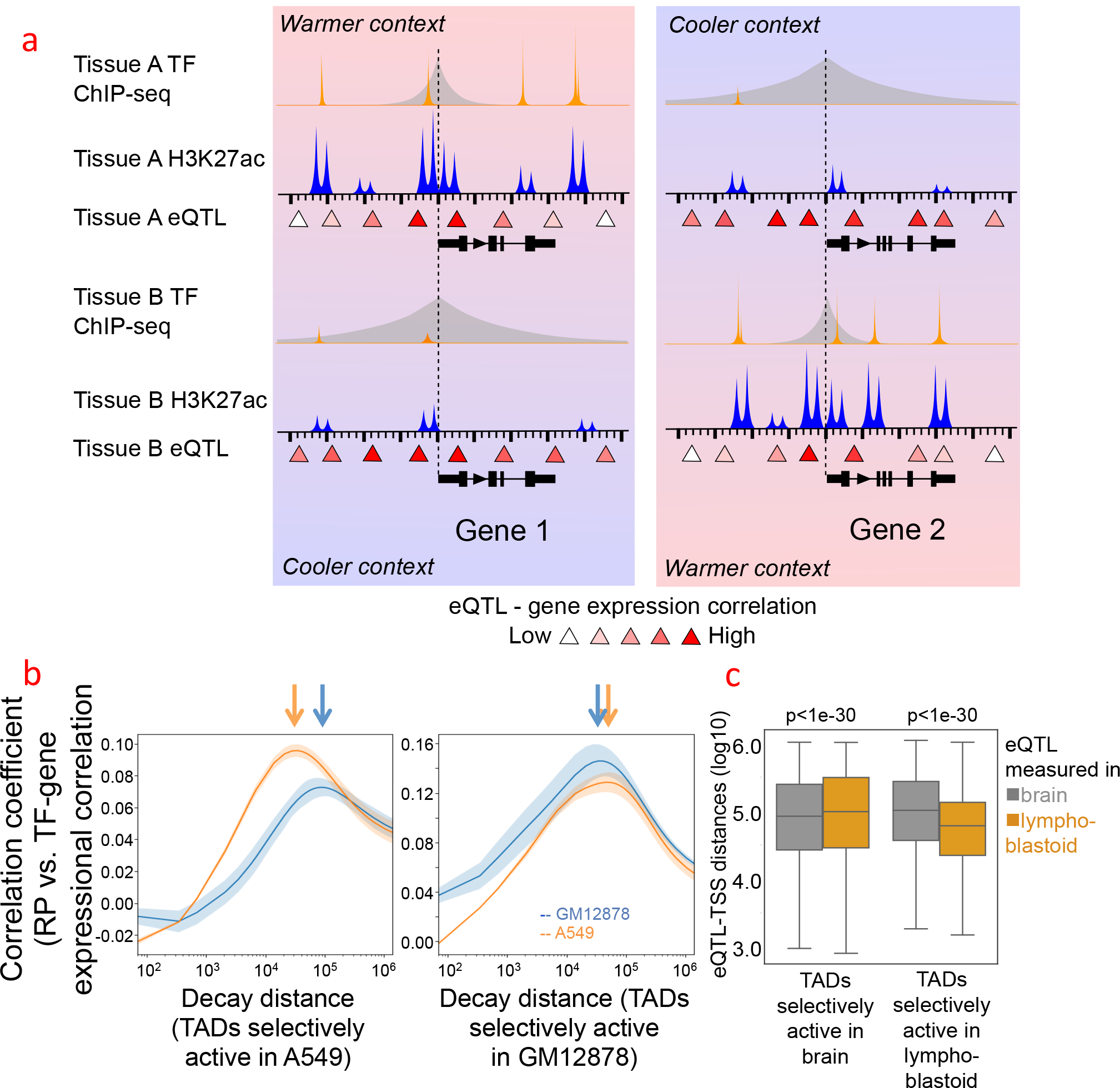
Regulatory decay distance changes with TAD activity and chromatin state. **(a)** Conceptual illustration of the idea that TF regulatory decay distances and eQTL-gene distances decrease as TADs become more active. **(b)** The regulatory decay distance of RELA decreases as TADs become active. RELA is expressed in both lymphoblastoid and lung, which possess distinct distributions of active TADs. In TADs more active in lung but less active in lymphoblastoid, the lung-specific RELA regulatory distance is shorter than lymphoblastoid-specific RELA regulatory distance (left). On the other hand, in TADs more active in lymphoblastoid but less active in lung, the lymphoblastoid-specific RELA regulatory distance is shorter than the lung-specific RELA regulatory distance (right). The lung-specific RELA regulatory decay distance is measured using RELA ChIP-seq in A549 lung cells and GTEx lung expression data, and the lymphoblastoid-specific RELA regulatory decay distance is measured using RELA ChIP-seq in GM12878 and GTEx lymphoblastoid expression data. **(c)** GTEx eQTL-TSS distances decrease in TADs that are active in the tissues in which the eQTL is measured. As (b), the distribution of GTEX eQTL-TSS distances measured in brain (gray) or lymphoblastoid (orange) were compared in brain-restricted or lymphoblastoid-restricted active TADs. Left: TADs more active in brain but less active in lymphoblastoid. Right: TADs more active in lymphoblastoid but less active in brain. The different log-transformed eQTL-TSS distances in individual group of TADs were compared using the two-sided Student t-test.

To further confirm this phenomenon, we examined the chromatin effects on regulatory decay distance using eQTL data from various tissues and cell types, including brain cortex, stomach, lymphoblastoid, and whole blood. We then identified the tissue-restricted TADs and determined whether the eQTLs within these TADs are closer to their associated genes in the tissues where these TADs are more active (Fig. 5a). For example, the eQTLs derived from brain cortex tissue tend to be closer to their associated genes in the TADs that are more active in brain. Likewise, lymphoblastoid eQTLs are closer to the associated genes in the TADs that are more active in lymphoblastoid cells (Fig. 5c). This is not a result of ascertainment bias because such bias would lead to the opposite result: there is greater statistical power to detect weak, distant, eQTL associations for highly expressed genes. In fact, this phenomenon holds true for most pairs of GTEx tissues with available H3K27ac ChIP-seq data we examined (Supplementary Fig. 4a), supporting our hypothesis that chromatin states of the TADs, in addition to genetic properties of the TADs, influence regulatory decay distances. More specifically, Δ* becomes shorter in the same TADs when the TADs become active.

### TAD-wise TF regulation model facilitates GWAS hit annotation and interpretation

Genome-wide association studies (GWAS) have mapped a large number of trait-associated single nucleotide polymorphisms (SNPs)^32^ in the non-coding regions of the genome, but the functional interpretation of these non-coding variants remains challenging. Several studies have shown that the GWAS SNPs are enriched in enhancer elements and DNase hypersensitive regions^33,34^. ENCODE^35^, ROADMAP^36^, GTEx^16^, and 4DN^37^ consortia have generated many useful data resources for interpreting GWAS SNPs^38^, but pinpointing causal variants and affected genes^38^ remains an open problem. Reports sometimes assign the causal gene as the nearest active gene to the GWAS hit^38^. The median distance between tag SNPs in the GWAS Catalog and the closest gene is ~11kb, which is far shorter than the regulatory decay distances of long-range TFs. Therefore, assigning GWAS SNPs to the nearest active genes might miss important targets. This is especially problematic if the GWAS SNPs are located in cold TADs and influence the expression of multiple distant genes in the same TAD. Indeed, among the GTEx eQTLs, GWAS SNPs are significantly enriched in cold TADs (Fig. 6a). Moreover, among the annotated eQTL-gene pairs in the GTEx eQTL data, GWAS SNPs interact with more genes (Fig. 6b) and with more distant genes (Fig. 6c) than non-GWAS SNPs. This underlies the importance of correctly modeling TF regulation decay distance in inferring causal genes of GWAS SNPs.

**Figure 6.**
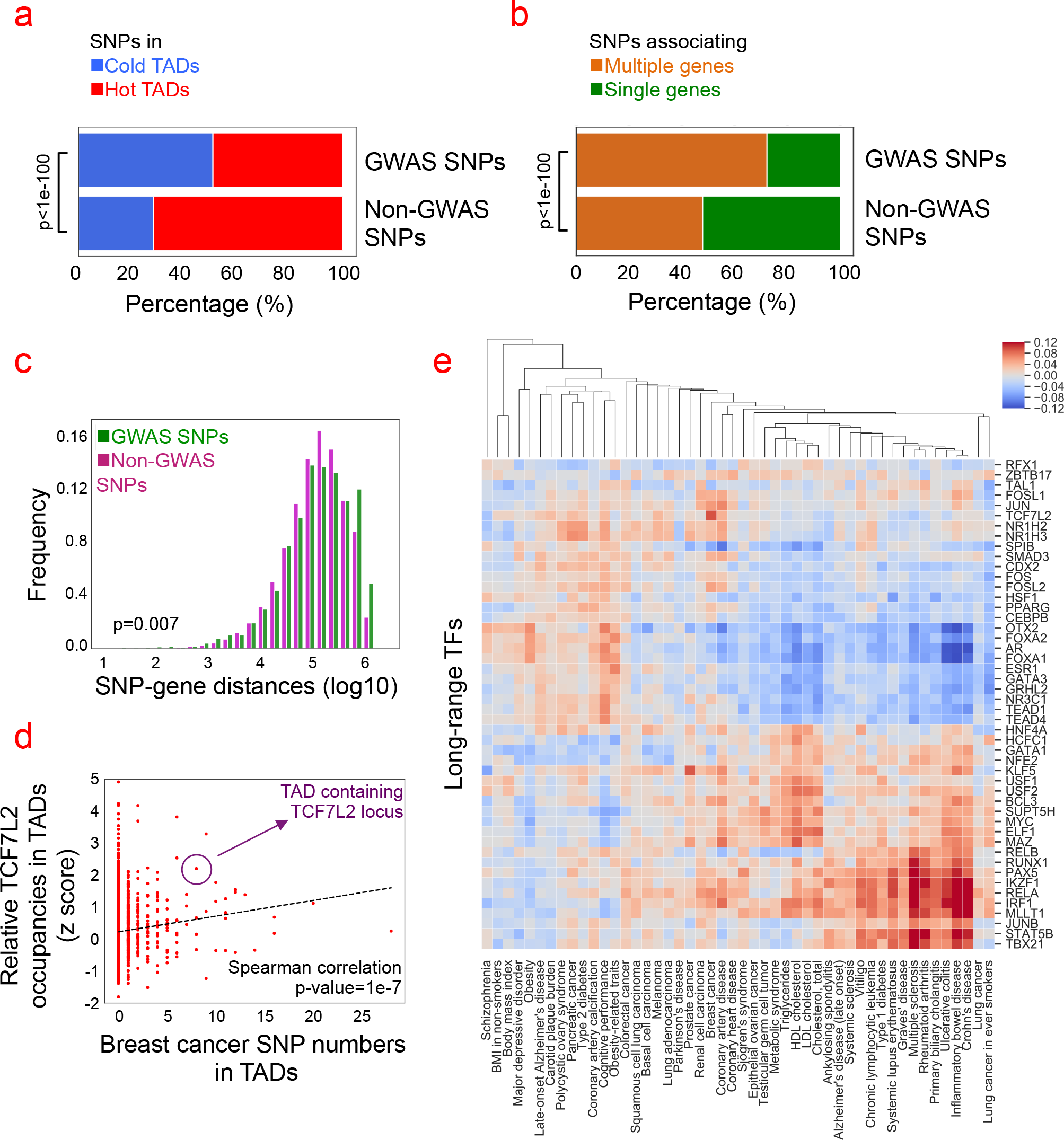
TAD-wise analysis of non-coding GWAS hits prioritizes relevant TFs. **(a)** Compared to non-GWAS SNPs, a higher proportion of GWAS SNPs are located in cold TADs (blue) than in hot TADs (red). The p-value calculated using the Fisher exact test. **(b-c)** Among annotated eQTL-gene pairs in the GTEx eQTL data, GWAS SNPs interact with more often with multiple genes (b) and with more distant genes (c) than non-GWAS SNPs. P-values were calculated using Fisher exact test (b) and two-sided Student t-test (c), respectively. **(d)** Breast cancer associated SNPs are enriched in TADs heavily bound by TCF7L2. Each dot represents one TAD. The x-axis represents the number of breast cancer associated SNPs, and the y-axis represents the relative TCF7L2 TF binding occupancies (z-score) in TADs. The purple circle indicates the TAD harboring the TCF7L2 gene locus. The p-value was calculated using the Spearman correlation between TCF7L2 occupancies and breast cancer SNPs numbers and the incomplete beta function. **(e)** Extension of calculation in (d) to all long-range TFs and GWAS traits, the heat map shows the enrichment or depletion of trait-associated SNPs in long-range TF target TADs. The x-axis represents traits annotated in the GWAS Catalog and the y-axis indicates long-range TFs. Each entry represents the Spearman correlation coefficient between relative TF occupancies in TADs and trait SNPs in TADs. Red squares are cases in which the indicated TFs and GWAS trait SNPs are co-localized in similar TADs. Blue squares are cases in which the indicated TFs tend to be depleted from TADs that contain GWAS SNPs of indicated trait. TFs and traits were further clustered using hierarchical clustering.

To demonstrate the use of these ideas to interpret GWAS SNPs, we used TCF7L2 as an example, which has been associated with breast cancer susceptibility^39,40^. TCF7L2 is a long-range TF, which based on our modeling tends to robustly regulate the genes located within its target TADs (Fig 2f). We compared the number of breast cancer SNPs and the number of TCF7L2 binding sites within each TAD and observed a significant positive correlation (Fig. 6d). The TAD containing the TCF7L2 locus itself (Fig. 6d) is particularly rich in TCF7L2 binding sites and in breast cancer SNPs. These results imply that the breast cancer SNPs not only regulate TCF7L2 expression, but also affect TCF7L2 downstream target genes in the TCF7L2 target TADs. Another example of a strong correlation between TF binding and GWAS SNP distribution is TBX21 in multiple sclerosis^41^ (Supplementary Fig. 5). This suggests that given SNPs associated with a particular disease, we could use this approach to implicate the long-range TFs involved in regulating the disease genes. Applying this approach to all the inferred long-range TFs we saw clusters of TFs to be associated with distinct groups of diseases, such as the previously known association between RUNX1 and autoimmune^42^ diseases, and between KLF5 and breast cancer^43^ (Fig. 6e). This analysis also revealed co-occurring diseases that have been reported in epidemiological studies, such as autoimmune diseases and chronic lymphocytic leukemia^44^, as well as type-2 diabetes and pancreatic cancer^45^. These findings demonstrate how a TAD-wise analysis of TF binding enrichment can be used to identify TFs germane to traits of interest in GWAS studies.

## Discussion

Despite intensive scientific investigation into the role of TFs in regulating metazoan gene expression, the mechanisms by which TFs regulate specific genes are still not well understood. In this study, we quantitatively modeled the ranges of genomic distance over which TFs regulate genes and the dependencies of these ranges on genomic and chromatin contexts. We systematically examined 3,406 TF ChIP-seq and 1,545 H3K27ac histone modification ChIP-seq datasets in combination with gene expression from 1,037 CCLE cancer cell lines and 48 GTEx tissues, and further analyzed eQTL and GWAS catalog data.

Our analyses revealed two distinct classes of TF with different ranges of regulatory influence, chromatin binding preferences, pioneer-like properties, and auto-regulatory behavior. Our results suggest that the binding sites of short-range TFs that are far away from gene TSSs, which account for the majority of their binding sites, usually don’t function to influence gene expression. Transcription is known to involve several distinct stages including recruitment of RNA polymerase II (PolII) to the promoter and release of the paused early elongation complex into productive elongation^46^. We speculate that short-range TFs might participate primarily in the Pol II recruitment stage while the long-range TFs might primarily influence elongation. A division in labor for transcriptional regulatory proteins has been reported for a small number of TFs and cofactors. For example, Bromodomain and ExtraTerminal (BET) proteins, mediator and the p-TEFb complexes are known to be important for productive transcriptional elongation^46^, and the TF Sp1 plays a known role in PolII preinitiation complex recruitment^47^. Our results suggest a division of labor applies to many DNA binding TFs. Recent observations that promoters can act like enhancers for other genes^48^ might depend on the recruitment of long-range TFs to enhancer-like promoters.

The genome is compartmentalized into topologically associating domains (TADs); some TADs are “cold” with low levels of activity in most cell types, while others are “hot” and highly active in most cell types. The known cases of genes being regulated by enhancers over extremely long genomic distances all occur in cold TADs, so we examined whether these observations were part of a general trend. Our analysis of TF ChIP-seq and eQTL data revealed a systematic trend in which the regulatory decay distances are longer in cold TADs than in hot ones. By measuring the regulatory distance in the same TADs under different chromatin states, we found that regulatory decay distances become shorter when chromatin becomes more active. As it is widely believed that enhancers regulate target genes through direct physical loop formation we explored Hi-C and Hi-ChIP data to determine if the interaction characteristics of hot and cold TADs could explain the distinct regulatory distances we observed, but they did not. It is possible that current interaction-detection technologies or data processing techniques do not capture all regulatory interactions on the relevant time and length scales or that non-looping mechanisms may mediate certain enhancer functions. The discordance between the interaction data and our decay distance analysis corresponds with recent high-resolution microscopy experiments on the regulation of SOX2^49^ and SHH^50^, which are regulated by certain distal enhancers without direct physical enhancer-promoter interaction.

Non-looping regulatory mechanisms may involve thermodynamically induced phase separation phenomena^51,52,53^. The compartments that have been observed in Hi-C data suggest that heterochromatin and euchromatin regions of the genome are spatially segregated^11^ and may form different nuclear structures such lamin-associated domains, nucleoli, nuclear speckles, PML bodies, and Cajal bodies^54,55^. In addition, altering the nuclear position of a locus has been observed to modify the expression of nearby genes^56,57^. Our analysis raises the possibility that TF-induced disruption of heterochromatin, by disruption of transcriptionally repressive zones or by nucleation of transcriptionally active ones, can lead to alterations in the regulatory microenvironment of genes and account for the chromatin state-dependent regulatory decay distances. Recent experimental work using a CRISPR-Cas9 based technology to induce phase-separated chromatin droplet formation at targeted genomic loci has shown that droplet formation in heterochromatin regions can cause large disruptions of heterochromatin domains^58^. The coordinated binding of pioneer factors to heterochromatin rich TADs could seed droplet nucleation points and the enhancer activity of these binding sites might be related to the release of a gene from a repressive environment. An alternative mechanism is facilitated tracking, in which enhancer-bound protein complexes move toward the promoter in a progressive, unidirectional fashion, while possibly remaining bound to the enhancer^59,60^. In principle, the presence of enhancers or promoters between the tracking enhancer and its target promoter may impede the tracking progress and would explain the shorter regulatory distances in hot domains.

Our modeling of regulatory decay distances is based on large TF cistrome data and gene expression cohorts of various cell lineages. Although the Cistrome DB contains most publicly available TF cistromes, and the CCLE and GTEx expression cohorts include many different cell and tissue types, we are far from covering all the TFs in all cellular contexts. Nonetheless, the high degree of consistency between analyses based on large amounts of orthogonal data provides solid support that the trends we observed are likely to reflect the general behavior of most TFs. These findings may help the interpretation of GWAS SNPs: significant SNPs in cold TADs can impact the expression of distant genes in the same TAD; significant SNPs in hot TADs, on the other hand, are less likely to be associated with (>100kb) distant genes. Some TADs are densely bound by certain long-range TFs, and the chromatin states of these TADs are dominated by these TFs. These TADs, which are “targets” of TFs, are predominantly of the “cold” type. Genes located in these target TADs usually have tissue-specific functions and the target TADs often harbor the TF gene itself, indicating TAD-wise auto-regulatory control of cell lineage specification. Integration of the heterogeneous distribution of long-range TFs and trait-associated SNPs in GWAS studies can help infer the TFs implicated in the manifested phenotype. Our findings highlight the importance of considering the TF type, genomic distance, and chromatin context in identifying TF targets and interpreting non-coding GWAS SNPs. Further work will be needed to understand the mechanistic basis of these TF-specific and context-dependent gene regulatory effects.

## Methods

### Key resource table

|  |  |
| --- | --- |
| GTEx | https://www.gtexportal.org/home/ |
| CTPR<sup>61</sup> | https://portals.broadinstitute.org/ctrp/ |
| Hi-C<sup>30</sup> | ftp://cooler.csail.mit.edu/coolers/hg19/ |
| ChIP-seq<sup>10</sup> | http://cistrome.org/db/#/ |
| GWAS<sup>32</sup> | https://www.ebi.ac.uk/gwas/home |
| Hi-ChIP<sup>62</sup> | https://www.ncbi.nlm.nih.gov/geo/query/acc.cgi?acc=GSE101498 |
| Linkage Disequilibrium block<sup>63,64</sup> | http://distild.jensenlab.org/download.html |
| TAD<sup>11</sup> | http://chromosome.sdsc.edu/mouse/hi-c/download.html |
| CAGE<sup>29</sup> | http://enhancer.binf.ku.dk/presets/ |

### TAD annotation

The TAD annotations were downloaded from http://chromosome.sdsc.edu/mouse/hi-c/download.html, and their coordinates were converted from hg18 to hg38 using liftOver software from UCSC: http://hgdownload.cse.ucsc.edu/goldenPath/hg18/liftOver/.

### Transcription factor ChIP-seq data processing and identification of enriched TADs

The raw sequence data of TF ChIPseq and H3K27ac ChIPseq were downloaded from Gene Expression Omnibus and processed through standard workflow of ChiLin^65^, consisting of quality control and peak calling using MACS^66^. For fair comparisons of TF occupancy distributions between samples, those TF ChIP-seq samples with less than 20,000 peaks were discarded, and for other TF ChIP-seq samples only the top 20,000 peaks based on peak intensities were included for downstream analysis.

For each TF ChIP-seq sample, we calculated the TF binding density in each TAD (Number of peaks / kb). Noticing that for all TFs this density is higher in the Hot TADs than the Cold TADs, we examined whether some TADs are more densely bound by certain TFs *relative to* others. To achieve this, for each TAD we calculated the mean and standard deviation of the TF binding density in the TAD across all TF samples. We then applied a z-score transformation to all TF ChIP-seq samples, and calculated the average z-score of a given TF to get the TAD-specific relative occupancies of the given TF compared to other TFs. For example, a TF *i* with a high z-score in a given TAD *j*, indicates that TAD *j* is more densely bound by TF *i* in comparison with other TFs. For each TF, we define TF target TADs as those with z-scores higher than 1.

### Regulatory potential model

In our model of transcription regulation by a given TF *i*, we assume that multiple TF *i* binding sites contribute additively to the regulation of a gene *j*. In this model, we modeled the effect of a single ChIP-seq peak *k* of TF *i* on gene *j* with an influence function that decays exponentially with the genomic distance between the TSS of gene *j* and peak *k*, *x*_*jk*_. The TF *i*-specific decay distance (Δ_*i*_) defines the half-life of the exponential decay function. The regulatory potential (RP), *R*_*i,j*_(Δ_*i*_), defines the total regulatory effect of TF *i* on gene *j* by summing all binding sites of TF *i* within range of the gene *j* (equation 1). Considering that the TAD boundary could insulate gene regulation, we summed up effects from the TF *i* peaks located with the same TAD as gene *j*.

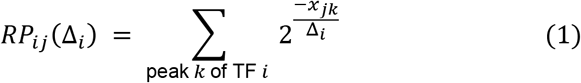

### Modeling regulatory distances of transcriptional factors

Assuming that 1) TFs are regulators of gene expression, and 2) TFs do not influence gene expression unless they are expressed, we can model transcriptional regulation of TF-*i* on expression of gene-*j* as

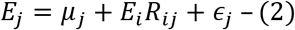

Where *E*_*j*_ is the expression of gene-*j*, *μ*_*j*_ is the basal expression of gene-*j*, *E*_*i*_ is the expression of TF-*i*, *R*_*ij*_ is the transcription regulatory effect of TF-*i* on of gene-*j*, and 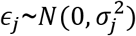.

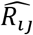, the maximum likelihood estimator of *R*_*ij*_, can be derived as:

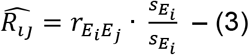

Where 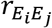 as the correlation coefficient between *E*_*i*_ and *E*_*j*_, while 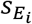 and 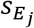 as the standard deviation of *E*_*i*_ and *E*_*j*_.

The transcription effect of TF-*i* on gene-*j*, *R*_*ij*_, can be modeled as:

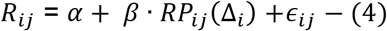

Where *RP*_*ij*_(Δ_*i*_) is the regulatory potential model of regulatory effect of TF-*i* on gene-*j*, defined as (1), and *ϵ*_*ji*_~*N*(0,σ^2^). With certain Δ_*i*_, *RP*_*ij*_(Δ_*i*_) is significantly correlated with 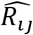 ( Supplementary Fig. 1a), suggesting our model *RP*_*ij*_(Δ_*i*_) is a good estimator of transcription regulatory effect of TF-*i* on of gene-*j*.

The parameters in equation (4) can be inferred using maximum likelihood estimations:

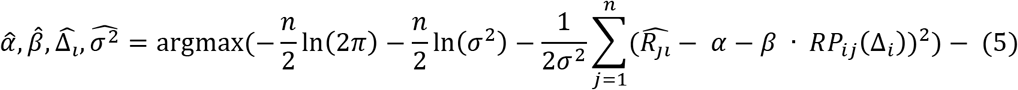

Where *n* is the number of genes.

To infer 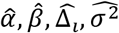, we first fixed Δ_*i*_, and 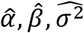 are:

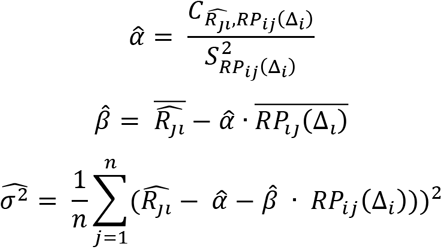

Then with fixed 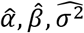,

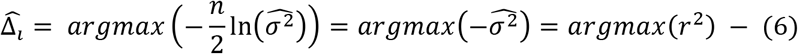

Where *r*, the Pearson correlation coefficient, is given by:

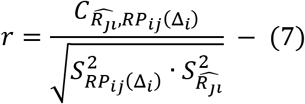

To estimate 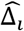, we iterated different Δ_*i*_ ranging from 100bp to 2000kb, and approximated 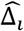 as the one that maximize *r*^2^. The standard deviation of 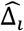 was estimated using bootstrapping. Specifically, with *n* genes, we re-sampled with replacement *n* samples from *n* genes, and re-estimated 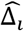. With *m* times re-sampling, we re-estimated 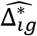, the *g*^*th*^ bootstrap estimator of 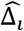.

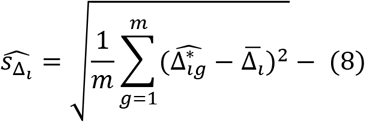

Where 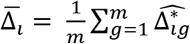

### Estimating regulatory distances of transcriptional factors using Cistrome TF ChIP-seq data and CCLE gene expression data

Based on equation (6), we can infer TF *i*-specific Δ_*i*_ by integrating TF ChIP-seq from Cistrome and gene expression data from CCLE database. Detailed steps are as follows:

1. We modeled *R*_*ij*_, the transcriptional regulatory effect of TF *i* on gene *j* as equation (1) using Cistrome TF ChIP-seq data and got *R*_*ij*_(Δ_*i*_).
2. As equation (3), 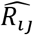, the maximum likelihood estimator of *R*_*ij*_ equals to the Pearson correlation 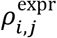 between the expression of TF *i* and gene *j*, using this normalized gene expression data. We downloaded CCLE gene expression data and normalized the log transformed RPKM values in different cell lines using quantile normalization, so that the distribution of gene expression values were the same in each cell line. Data were then normalized gene-wise by mean centering.
3. As equation (6), 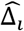 can be inferred as the Δ that gives rise to highest 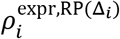. Therefore, we repeated the above step 1-2 with Δ_*i*_ ranging from 100bp to 2000kb, and defined 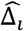, the “regulatory decay distance” of TF *i* as:

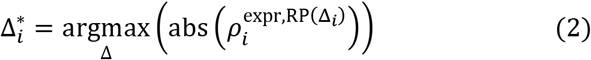 Specifically, we inferred the optimal Δ using the following steps:

a. We defined qualified TF ChIP-seq samples as those producing a maximum absolute value of 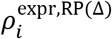 greater than 0.1.
b. If there were more than 2 qualified TF *i* ChIP-seq samples, we averaged the 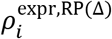 for each Δ. If the maximum average correlation is larger than 0.1, we chose the Δ that gives rise to the maximum average correlation coefficient as the “regulatory decay distance” of TF *i*.

### Modeling dynamic regulatory distances of transcriptional factors

For the investigation of the regulatory decay distance dependence on chromatin context, we proceeded as follows:

1. To infer TAD type-specific (e.g. Cold TADs or Hot TADs) regulatory decay distances, in step 3 of “Modeling regulatory distances of transcriptional factor using CCLE gene expression data” we included only genes located within the designated TAD type (e.g. Cold TADs or Hot TADs). Other steps remained the same.
2. To investigate whether the regulatory decay distances depend on chromatin status in addition to genomic features, we focused on TADs that have distinct activity status levels in different lineages. Take lung and lymphoblastoid tissues, for example. We first computed the lung H3K27ac level for each TAD by averaging the TAD H3K27ac mean values over all H3K27ac ChIP-seq samples associated with the lung cluster. The lymphoblastoid H3K27ac TAD levels were calculated in a similar way. We defined the lung-specific TAD clusters as those with significantly (p-value<1e-10) higher H3K27ac levels in lung compared with H3K27ac ChIP-seq produced from lymphoblastoid. The lymphoblastoid-specific TAD clusters were defined in a similar way. For the RELA analysis we used GTEx lung expression data and RELA ChIP-seq samples in A549 to compute the lung-specific regulatory decay distance. RELA ChIP-seq in lymphoblastoid and GTEx lymphoblastoid expression data was used to compute the lymphoblastoid-specific regulatory decay distance.

### H3K27ac clustering

The raw sequence data of TF ChIP-seq and H3K27ac ChIP-seq were downloaded from Gene Expression Omnibus and processed through standard workflow of Chilin^65^ with peak calling using MACS^66^ and the processed data is accessible through the Cistrome Data Browser. We calculated the average H3K27ac occupancies in each TAD using bigWigAverageOverBed, and further normalized each sample using z-transformation. We then clustered the H3K27ac CHIP-seq samples and TAD as 10 clusters, both using hierarchical clustering. We defined the TAD cluster with the weakest and strongest H3K27ac signals as Cold TADs and Hot TADs, respectively.

### Pioneer-like factors identification

Pioneer factors can access closed chromatin and facilitate the binding of other TFs to the accessible genomic loci. To probe whether a TF such as TEAD1, for example, possesses pioneer-like factor properties, we investigated whether there is a gain in TEAD1 motif enrichment in peaks of non-TEAD1 TF ChIP-seq samples in cell lines with high levels of TEAD1 expression. We processed as following steps:

1. Calculate the TEAD1 motif enrichment in peaks of all non-TEAD1 TF ChIP-seq samples using Homer^67^. TEAD1 motif enrichment represents the ratio of the proportion of non-TEAD1 ChIP-seq peaks with the TEAD1 motif to the proportion of background sequences with the TEAD1 motif.
2. Average the TEAD1 motif enrichment for redundant TF-cell pairs.
3. Define the median TEAD1 enrichment across all non-TEAD1 TF ChIP-seq samples in each cell line *i* as *y*_*i*_ and define *x*_*i*_ as the level of TEAD1 gene expression in that cell line. Defining *ӯ* as the mean of *y*_*i*_ across all cell lines, we further derived 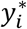 as follows:

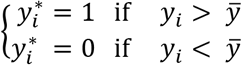
4. Perform logistic regression 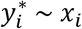, and define TEAD1 as “pioneer-factor like” if the logistic regression p-value is smaller than 0.01.

### GTEx eQTL analysis

We downloaded eQTL loci across multiple tissues from the Genotype-Tissue Expression (GTEx) database (accession phs000424.v7.p2), remapped the coordinates from GRCh37/hg19 to GRCh38/hg38, and filtered out eQTL-TSS pairs with p-values smaller than 1e-5. To investigate whether eQTL-TSS pairs in Cold TADs have longer distances than those in Hot TADs, we selected those eQTL-TSS pairs located in Cold and Hot TADs, and then compared their log-transformed eQTL-TSS distances using Student’s t-test. To further examine whether the eQTL-TSS distances become shorter as chromatin becomes more active, we focused on TADs that have distinct activities in different lineages. Take brain cortex and lymphoblastoid for instance, we compared their H3K27ac level using Cistrome and defined brain-specific active TADs as those with significantly (p-value<1e-10) higher H3K27ac level in brain cortex compared with lymphoblastoid. Lymphoblastoid-specific active TADs were defined in the same way. Further for eQTL-TSS pairs located in brain-specific or lymphoblastoid-specific active TADs, we compared the log-transformed distances measured in brain cortex and lymphoblastoid using GTEx data.

For more comprehensive comparisons, we compared all possible pairs from whole blood, stomach, lymphoblastoid, and brain cortex. We defined those pairs with eQTL-TSS distances longer in tissue-specific active TADs as non-significant.

### Hi-C analysis

The Hi-C data in cool format was downloaded from ftp://cooler.csail.mit.edu/coolers/hg19/. We mapped contact pairs to cold and hot TADs, respectively, and compared the decreasing rate of contact frequencies with distances. Specifically, for given genomic distance (*d*), we calculated the average contact frequencies (*F*):

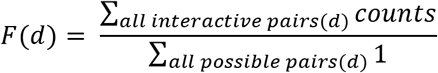

### GWAS and GTEx eQTL join analysis

We classified the SNPs in GTEx eQTL as those that are annotated in GWAS catalog (“GWAS SNPs”) and others (“Non-GWAS SNPs”).

### GWAS SNP enrichment analysis

We mapped the GWAS catalog SNPs to TADs, and then for each long-range TF vs. GWAS trait pair we calculated the Spearman correlation coefficients of SNPs numbers and relative TF enrichment on TADs. We plotted the diseases/traits – TF heatmap with correlation coefficients, and clustered the diseases/traits and TFs using hierarchical clustering.

## AUTHOR CONTRIBUTIONS

C.-H.C. and C.M. conceived the project. C.-H.C performed the analyses. C.M and X.S.L supervised the whole study and wrote the manuscript with C.-H.C. with the help of all other authors.

## ACKNOWLEDGEMENTS

We thank Drs. Leonid Mirny, Bing Ren, and Fulai Jin for helpful discussion during manuscript preparation. We also acknowledge the following funding sources for supporting our work: National Key Research and Development Program of China 2017YFC0908500 to X.S.L., and NIH grant U24 HG009446 and U01 CA180980 to X.S.L.

## References

1. Banerji, J., Olson, L. & Schaffner, W. A lymphocyte-specific cellular enhancer is located downstream of the joining region in immunoglobulin heavy chain genes. Cell 33, 729–40 (1983).

2. Gillies, S. D., Morrison, S. L., Oi, V. T. & Tonegawa, S. A tissue-specific transcription enhancer element is located in the major intron of a rearranged immunoglobulin heavy chain gene. Cell 33, 717–28 (1983).

3. Long, H. K., Prescott, S. L. & Wysocka, J. Ever-Changing Landscapes: Transcriptional Enhancers in Development and Evolution. Cell 167, 1170–1187 (2016).

4. Fukaya, T., Lim, B. & Levine, M. Enhancer Control of Transcriptional Bursting. Cell 166, 358–368 (2016).

5. Ouyang, Z., Zhou, Q. & Wong, W. H. ChIP-Seq of transcription factors predicts absolute and differential gene expression in embryonic stem cells. Proc. Natl. Acad. Sci. U. S. A. 106, 21521–6 (2009).

6. Robertson, G. et al. Genome-wide profiles of STAT1 DNA association using chromatin immunoprecipitation and massively parallel sequencing. Nat. Methods 4, 651–657 (2007).

7. Johnson, D. S., Mortazavi, A., Myers, R. M. & Wold, B. Genome-Wide Mapping of in Vivo Protein-DNA Interactions. Science (80-.). 316, 1497–1502 (2007).

8. Mikkelsen, T. S. et al. Genome-wide maps of chromatin state in pluripotent and lineage-committed cells. Nature 448, 553–60 (2007).

9. Barski, A. et al. High-Resolution Profiling of Histone Methylations in the Human Genome. Cell 129, 823–837 (2007).

10. Mei, S. et al. Cistrome Data Browser: a data portal for ChIP-Seq and chromatin accessibility data in human and mouse. Nucleic Acids Res. 45, D658–D662 (2017).

11. Dixon, J. R. et al. Topological domains in mammalian genomes identified by analysis of chromatin interactions. Nature 485, 376–380 (2012).

12. Pope, B. D. et al. Topologically associating domains are stable units of replication-timing regulation. Nature 515, 402–405 (2014).

13. Long, H. K., Prescott, S. L. & Wysocka, J. Leading Edge Review Ever-Changing Landscapes: Transcriptional Enhancers in Development and Evolution. Cell 167, 1170–1187 (2016).

14. Rada-Iglesias, A. et al. A unique chromatin signature uncovers early developmental enhancers in humans. Nature 470, 279–283 (2011).

15. Barretina, J. et al. The Cancer Cell Line Encyclopedia enables predictive modelling of anticancer drug sensitivity. Nature 483, 603–7 (2012).

16. Lonsdale, J. et al. The Genotype-Tissue Expression (GTEx) project. Nat. Genet. 45, 580–585 (2013).

17. Wang, S. et al. Target analysis by integration of transcriptome and ChIP-seq data with BETA. Nat. Protoc. 8, 2502–2515 (2013).

18. Flavahan, W. A. et al. Insulator dysfunction and oncogene activation in IDH mutant gliomas. Nature 529, 110–4 (2016).

19. Nora, E. P. et al. Spatial partitioning of the regulatory landscape of the X-inactivation centre. Nature 485, 381–385 (2012).

20. Duboule, D. The rise and fall of Hox gene clusters. Development 134, 2549–60 (2007).

21. Dekker, J. GC- and AT-rich chromatin domains differ in conformation and histone modification status and are differentially modulated by Rpd3p. Genome Biol. 8, R116 (2007).

22. Zaret, K. S. & Carroll, J. S. Pioneer transcription factors: establishing competence for gene expression. Genes Dev. 25, 2227–41 (2011).

23. Jacobs, J. et al. The transcription factor Grainy head primes epithelial enhancers for spatiotemporal activation by displacing nucleosomes. Nat. Genet. 50, 1011–1020 (2018).

24. Palani, S. & Sarkar, C. A. Positive receptor feedback during lineage commitment can generate ultrasensitivity to ligand and confer robustness to a bistable switch. Biophys. J. 95, 1575–89 (2008).

25. Crews, S. T. & Pearson, J. C. Transcriptional autoregulation in development. Curr. Biol. 19, R241–6 (2009).

26. Isoda, T. et al. Non-coding Transcription Instructs Chromatin Folding and Compartmentalization to Dictate Enhancer-Promoter Communication and T Cell Fate. Cell 171, 103–119.e18 (2017).

27. Ohmura, S. et al. Lineage-affiliated transcription factors bind the Gata3 Tce1 enhancer to mediate lineage-specific programs. J. Clin. Invest. 126, 865–878 (2016).

28. Osterwalder, M. et al. Enhancer redundancy provides phenotypic robustness in mammalian development. Nature 554, 239–243 (2018).

29. Andersson, R. et al. An atlas of active enhancers across human cell types and tissues. Nature 507, 455–461 (2014).

30. Dixon, J. R. et al. Chromatin architecture reorganization during stem cell differentiation. Nature 518, 331–336 (2015).

31. Lieberman-Aiden, E. et al. Comprehensive mapping of long-range interactions reveals folding principles of the human genome. Science 326, 289–93 (2009).

32. MacArthur, J. et al. The new NHGRI-EBI Catalog of published genome-wide association studies (GWAS Catalog). Nucleic Acids Res. 45, D896–D901 (2017).

33. Trynka, G. et al. Chromatin marks identify critical cell types for fine mapping complex trait variants. Nat. Genet. 45, 124–130 (2013).

34. Schaub, M. A., Boyle, A. P., Kundaje, A., Batzoglou, S. & Snyder, M. Linking disease associations with regulatory information in the human genome. Genome Res. 22, 1748–59 (2012).

35. ENCODE Project Consortium, T. E. P. An integrated encyclopedia of DNA elements in the human genome. Nature 489, 57–74 (2012).

36. Bernstein, B. E. et al. The NIH Roadmap Epigenomics Mapping Consortium. Nat. Biotechnol. 28, 1045–1048 (2010).

37. Dekker, J. et al. The 4D nucleome project. Nature 549, 219–226 (2017).

38. Edwards, S. L., Beesley, J., French, J. D. & Dunning, A. M. Beyond GWASs: illuminating the dark road from association to function. Am. J. Hum. Genet. 93, 779–97 (2013).

39. Lu, X. P., Hu, G. N., Du, J. Q. & Li, H. Q. TCF7L2 gene polymorphisms and susceptibility to breast cancer: a meta-analysis. Genet. Mol. Res. 14, 2860–2867 (2015).

40. Burwinkel, B. et al. Transcription factor 7-like 2 (TCF7L2) variant is associated with familial breast cancer risk: a case-control study. BMC Cancer 6, 268 (2006).

41. Patsopoulos, N. A. et al. Genome-wide meta-analysis identifies novel multiple sclerosis susceptibility loci. Ann. Neurol. 70, 897–912 (2011).

42. Alarcón-Riquelme, M. E. Role of RUNX in autoimmune diseases linking rheumatoid arthritis, psoriasis and lupus. Arthritis Res. Ther. 6, 169–73 (2004).

43. Jia, L. et al. KLF5 promotes breast cancer proliferation, migration and invasion in part by upregulating the transcription of TNFAIP2. Oncogene 35, 2040–2051 (2016).

44. Hodgson, K., Ferrer, G., Montserrat, E. & Moreno, C. Chronic lymphocytic leukemia and autoimmunity: a systematic review. Haematologica 96, 752–61 (2011).

45. Li, D. Diabetes and pancreatic cancer. Mol. Carcinog. 51, 64–74 (2012).

46. Adelman, K. & Lis, J. T. Promoter-proximal pausing of RNA polymerase II: emerging roles in metazoans. Nat. Rev. Genet. 13, 720–731 (2012).

47. Dorris, D. R. & Struhl, K. Artificial Recruitment of TFIID, but Not RNA Polymerase II Holoenzyme, Activates Transcription in Mammalian Cells. 20, (2000).

48. Andersson, R., Sandelin, A. & Danko, C. G. A unified architecture of transcriptional regulatory elements. Trends Genet. 31, 426–433 (2015).

49. Alexander, J. M., Guan, J., Huang, B., Lomvardas, S. & Weiner, O. D. Live-Cell Imaging Reveals Enhancer-dependent Sox2 Transcription in the Absence of Enhancer Proximity. bioRxiv 409672 (2018). doi:10.1101/409672

50. Benabdallah, N. S. et al. PARP mediated chromatin unfolding is coupled to long-range enhancer activation. bioRxiv 155325 (2017). doi:10.1101/155325

51. Strom, A. R. et al. Phase separation drives heterochromatin domain formation. Nature 547, 241–245 (2017).

52. Larson, A. G. et al. Liquid droplet formation by HP1α suggests a role for phase separation in heterochromatin. Nature 547, 236–240 (2017).

53. Savić, N. et al. lncRNA maturation to initiate heterochromatin formation in the nucleolus is required for exit from pluripotency in ESCs. Cell Stem Cell 15, 720–34 (2014).

54. van Steensel, B. & Belmont, A. S. Lamina-Associated Domains: Links with Chromosome Architecture, Heterochromatin, and Gene Repression. Cell 169, 780–791 (2017).

55. Quinodoz, S. A. et al. Higher-Order Inter-chromosomal Hubs Shape 3D Genome Organization in the Nucleus. Cell (2018). doi:10.1016/j.cell.2018.05.024

56. Finlan, L. E. et al. Recruitment to the Nuclear Periphery Can Alter Expression of Genes in Human Cells. PLoS Genet. 4, e1000039 (2008).

57. Andrulis, E. D., Neiman, A. M., Zappulla, D. C. & Sternglanz, R. Perinuclear localization of chromatin facilitates transcriptional silencing. Nature 394, 592–595 (1998).

58. Shin, Y. et al. Liquid Nuclear Condensates Mechanically Sense and Restructure the Genome. Cell 175, 1481–1491.e13 (2018).

59. Blackwood, E. M. & Kadonaga, J. T. Going the distance: a current view of enhancer action. Science 281, 60–3 (1998).

60. Vernimmen, D. & Bickmore, W. A. The Hierarchy of Transcriptional Activation: From Enhancer to Promoter. Trends Genet. 31, 696–708 (2015).

61. Rees, M. G. et al. Correlating chemical sensitivity and basal gene expression reveals mechanism of action. Nat. Chem. Biol. 12, 109–116 (2016).

62. Mumbach, M. R. et al. Enhancer connectome in primary human cells identifies target genes of disease-associated DNA elements. Nat. Genet. 49, 1602–1612 (2017).

63. Palleja, A., Horn, H., Eliasson, S. & Jensen, L. J. DistiLD Database: diseases and traits in linkage disequilibrium blocks. Nucleic Acids Res. 40, D1036–D1040 (2012).

64. Gibbs, R. A. et al. The International HapMap Project. Nature 426, 789–796 (2003).

65. Qin, Q. et al. ChiLin: a comprehensive ChIP-seq and DNase-seq quality control and analysis pipeline. BMC Bioinformatics 17, 404 (2016).

66. Zhang, Y. et al. Model-based Analysis of ChIP-Seq (MACS). Genome Biol. 9, R137 (2008).

67. Heinz, S. et al. Simple combinations of lineage-determining transcription factors prime cis-regulatory elements required for macrophage and B cell identities. Mol. Cell 38, 576–89 (2010).

